# GlueFinder: A Data-Driven Framework for the Rational Discovery of Molecular Glues

**DOI:** 10.1101/2025.10.17.683126

**Authors:** Jeffrey Skolnick, Bharath Srinivasan, Hongyi Zhou

**Affiliations:** Center for the Study of Systems Biology, Georgia Institute of Technology, 950 Atlantic Dr NW, Atlanta, GA 30332, USA; School of Pharmacy and Life Sciences, Robert Gordon University, Garthdee House, Garthdee Rd, Aberdeen AB10 7AQ, UK; Department of Chemistry, Stony Brook University, Stony Brook NY, 11794-3400, USA; Center for the Advanced Study of Drug Action, Stony Brook University, Stony Brook NY, 11794-3400, USA; Cancer Research Horizons, Cancer Research UK, 2 Redman Place. London. E20 1JQ, UK

**Keywords:** Molecular glue, degraders, induced proximity, E3 ligase, rational design

## Abstract

Molecular glues can drive targeted protein degradation by stabilizing ternary complexes between proteins of interest and E3 ubiquitin ligases, but rational design has lagged due to limited rules for interface recognition and an overreliance on a few ligases (e.g., VHL or Cereblon). We introduce GlueFinder, a systematic, unbiased platform that leverages structural bioinformatics to mine the Protein Data Bank for ligand binding pockets adjacent to the protein interface which are ligandable sites near protein–protein interfaces that can nucleate glue-mediated complex formation. After validating its performance on a benchmark of experimentally solved dimeric structures with known and predicted glues, we applied GlueFinder to three therapeutically important targets, EGFR, HER2, and KRAS, and predicted candidate glues that recruit 24, 111, and 148 distinct E3 ligases to these targets, respectively. We further demonstrate that GlueFinder can promote the formation of non-native EGFR complexes, possibly enabling ternary assemblies that would not form on their own. Together, these results establish a general, computation-guided strategy for molecular glue discovery that decouples design from legacy degrader scaffolds and specific ligase dependencies, expands the usable E3 ligase repertoire, and enables rational targeting of interfacial binding pockets. GlueFinder thus broadens both the scope and precision of targeted protein degradation and moves the field toward mechanism-driven, systematic glue development across diverse therapeutic contexts.

**Significance:** Molecular glues are small molecules that destroy disease-causing proteins by helping them attach to cellular disposal machinery. However, current glue discovery depends on a few known ligases and lacks clear design rules. GlueFinder provides a general, computation-guided method that scans protein structures to find pockets near interfaces where glues can act. Applied to key targets such as EGFR, HER2, and KRAS, it predicts glues that connect many different ligases and even create new protein complexes. By revealing where and how glues can work, GlueFinder expands the range of degradable proteins and accelerates the rational design of future therapeutics.

## Introduction

Molecular glues are an emerging class of small molecules that act by stabilizing protein–protein interactions (PPIs), enabling novel therapeutic mechanisms that diverge significantly from conventional small-molecule inhibitors (1). Unlike traditional drugs, which typically inhibit the activity of a single target protein by occupying active or allosteric sites, as shown in Figure 1, molecular glues function by *inducing or stabilizing* interactions between two proteins, often between a target protein and a cellular effector, thereby reprogramming protein function or fate. This mechanism of action also sets molecular glues apart from proteolysis-targeting chimeras (PROTACs), which rely on bifunctional molecules to physically tether a target protein to an E3 ubiquitin ligase (2). While PROTACs require defined binding motifs for both the target and the ligase, molecular glues operate through subtle surface complementarity and induced proximity, often modifying interactions that are otherwise transient or non-existent (see Figure 1).

**Figure 1.**
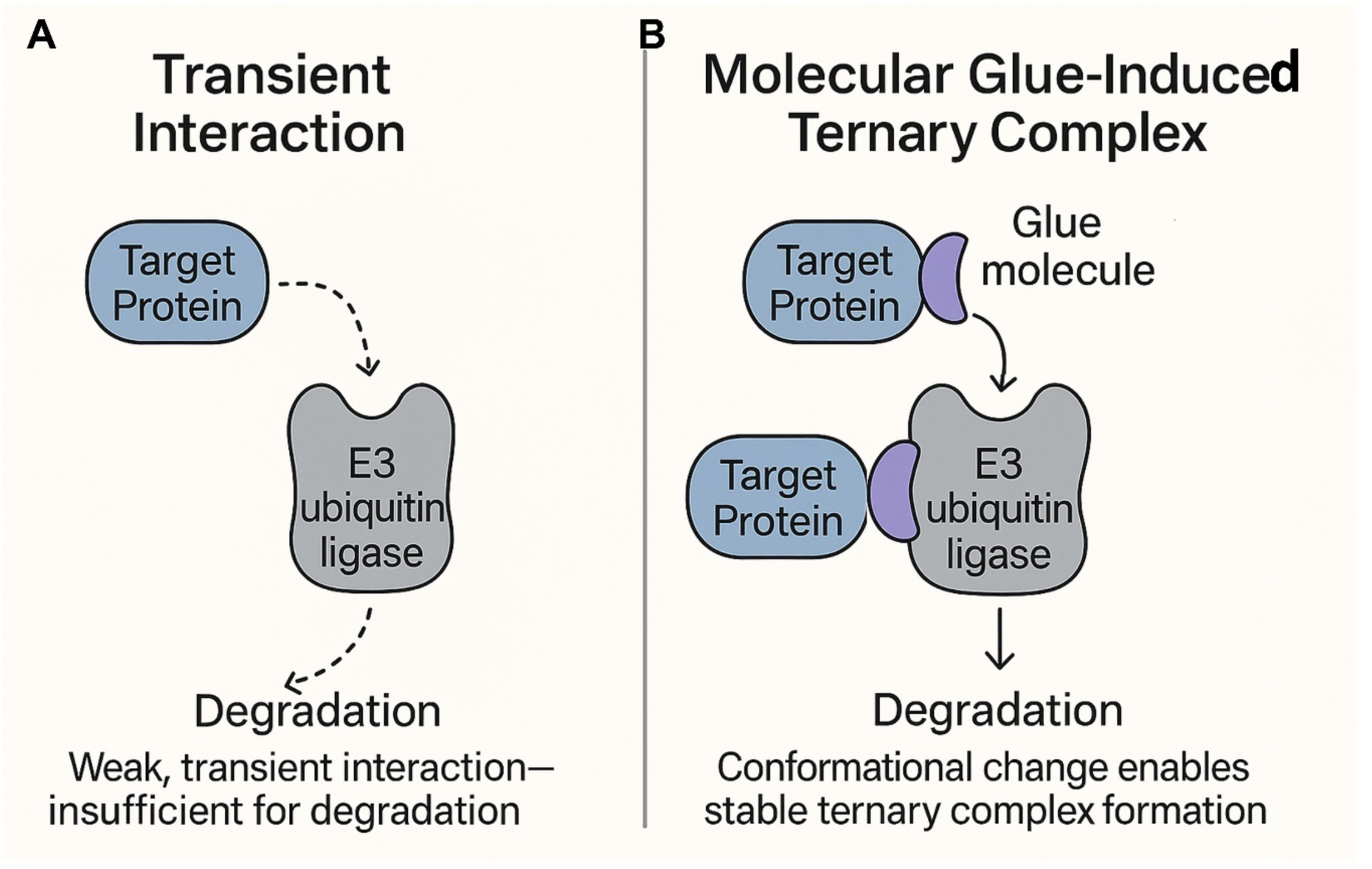
Schematic representation of mechanism of action for molecular glues. In the cellular milieu, protein-protein interactions are often transient. Molecular glues, as emphasized by the example of molecular glue degraders shown here, stabilize these interactions and make them constitutive, resulting in rewiring the cellular metabolic machinery and controls.

Molecular glues offer several compelling advantages. Their small size facilitates better cell permeability, blood-brain barrier (BBB) penetrance and favorable pharmacokinetic properties compared to larger bifunctional molecules like PROTACs (1, 3). They can also access previously “undruggable” proteins that lack suitable binding pockets for traditional inhibitors or induce protein-protein complexes when the complex would be unstable in the absence of the molecular glue. Moreover, by leveraging endogenous degradation or signaling pathways, molecular glues can exert catalytic effects, modulating protein function even at sub-stoichiometric doses (4). However, their development presents significant challenges. Most molecular glues have been discovered serendipitously, often through phenotypic screening rather than rational design (3). The precise molecular determinants that govern glue-induced ternary complex formation and functional outcomes remain incompletely understood, limiting the ability to generalize from known scaffolds or targets.

The therapeutic promise of molecular glues is becoming increasingly evident, particularly in oncology, where they have shown effectiveness in degrading oncogenic transcription factors and other difficult-to-target proteins (2). For instance, thalidomide and its analogues (IMiDs) have demonstrated the power of molecular glues to modulate substrate recognition by E3 ligases such as Cereblon (CRBN), leading to targeted degradation of neosubstrates like IKZF1/3 (5). Beyond cancer, molecular glues are being explored for neurodegenerative diseases, immunological disorders, and viral infections, expanding their reach as a broad therapeutic modality (1, 6).

A notable subclass of these molecules, molecular glue degraders, function by enhancing or enforcing interactions between E3 ubiquitin ligases and target proteins of interest (POI), leading to ubiquitin-mediated proteasomal degradation (Figure 1). These degraders essentially transform weak or transient PPIs into stable complexes, redirecting the cellular degradation machinery to specific targets (7). By reprogramming the substrate specificity of E3 ligases, they offer a powerful strategy for selectively depleting disease-causing proteins, including those lacking enzymatic activity or conventional drug-binding sites. A very prominent class of molecular glue degraders are the immunomodulatory imide drugs (IMiDs) such as thalidomide and its derivatives, lenalidomide (Revlimid) and pomalidomide (Pomalyst) (5), see Figure 2A. These compounds bind to the Cereblon (CRBN) protein, which is part of a larger ubiquitin ligase complex. This binding creates a new interaction interface, effectively acting as a “glue” to recruit specific target proteins, such as the transcription factors IKZF1 and IKZF3, to the CRBN-E3 ligase complex. Depending on the situation at hand, they can act as a “bridge compound” (see Figure 3 below) such as between CRBN and SAL. In other cases, such as the case of pomalidomide binding to CRBN and IKAROS, the CRBN domain in which pomalidomide is bound, has a RMSD of 1.12 Å to the monomeric structure found in 5ank; however, it forms is an interface adjacent ligand with IKAROS. In a number of other cases, the structure alignment of monomeric CRBN (PDB id: 8rq8A (8)) to its structure in found in 4Ci1B (9), 8oizB (10), 8g66B (11), 5fqdB (12), 5hxbC (13), 6h0gB (14), 8u16A (15), and 9tnpB (16) dimers are all below 2.1Å. 8rq8A (a monomer) lacks an inserted helical hairpin that interacts with the partner protein. The structure of 8tnp is typical where CBRN interacts with DNA damage protein 1 and pomalidomide is an IAP binding ligand (see Figure S1A.8tnp and the superposition onto the monomeric 8rq8A, Figure S1B. 8rq8A.8tnp). The structural alignment of CRBN in 8rq8A and 6hf0B is 3.44Å (see Figure S2. 8rq8A.6hf0B.cartoon) where the pomalidomide binding domain opens and detaches from the remainder of the protein to interact with IKAROS. However, pomalidomide is still an IAP binding ligand. Thus, the algorithms described below could predict the binding of CRBN with IKAROS even if they rely on a structure prediction algorithm such as AlphaFold 2 (AF2), AlphaFold 3 (AF3) or its variants to predict the quaternary structure.

**Figure 2.**
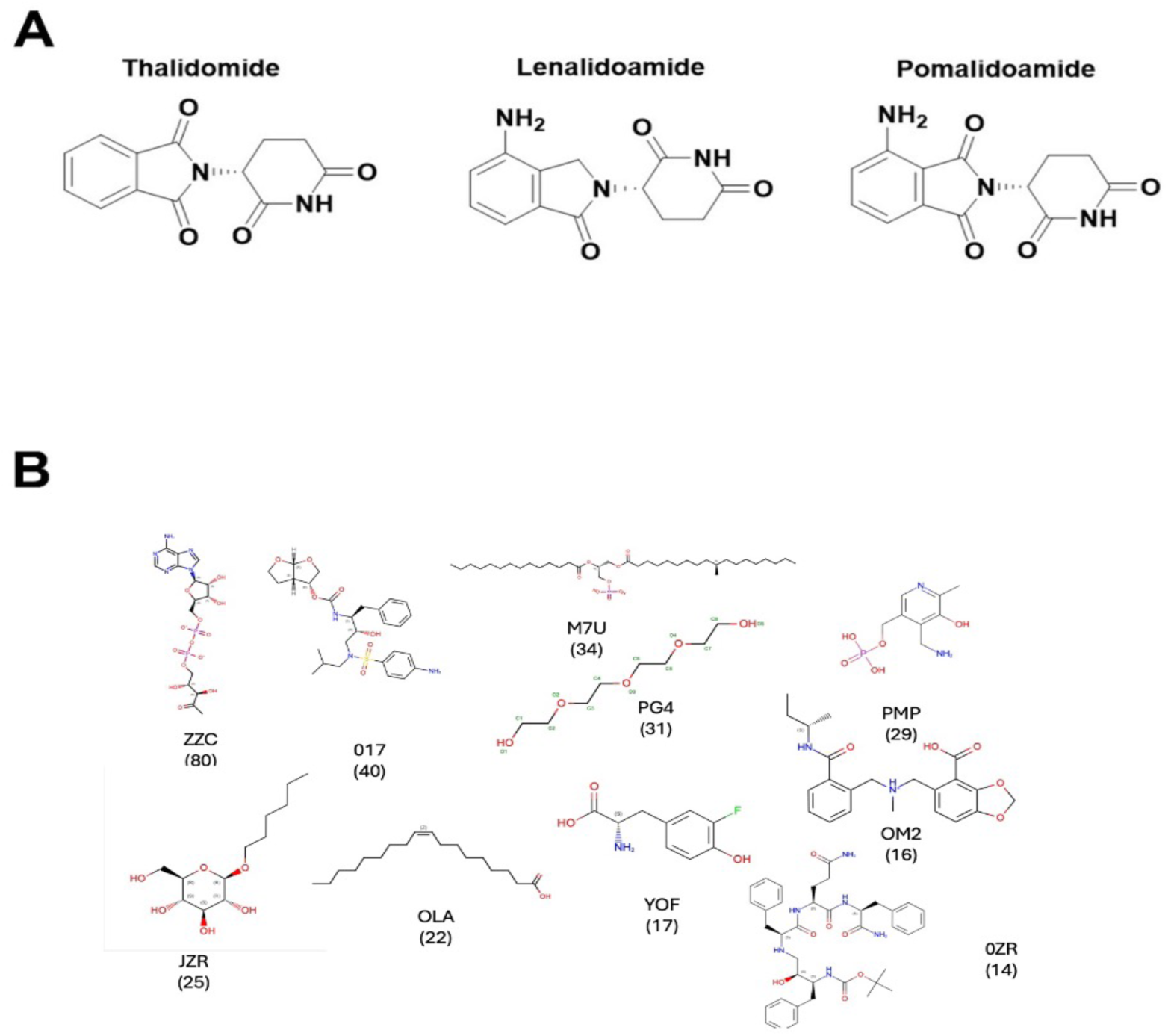
**A** IMiD class of molecular glue degraders thalidomide and its derivatives, lenalidomide (Revlimid) and pomalidomide (Pomalyst). These recruit CRBN E3 ligase to degrade the protein targets of interest. **B.** Cluster centroids of the top 10 clusters of interface adjacent ligands found in the PDB: ZZC ([(2r,3s,4r,5r)-5-(6-aminopurin-9-yl)-3,4-dihydroxy-oxolan-2-yl]methyl [[(2r,3r)-2,3-dihydroxy-4-oxo-pentoxy]-oxido-phosphoryl] phosphate); 017 (3R,3S,6R)-hexahydrofuro[2,3-b]furan-3-yl(1s,2r)-3-[[(4-aminophenyl) sulfonyl] (isobutyl)amino]-1-benzyl-2-hydroxypropylcarbamate); M7U (2R)-2-(hexadecanoyloxy)- 3([(10r)-10-methyloctadecanoyl]oxy}propyl phosphate; PG4 (tetraethylene glycol); PMP (4’-deoxy-4’-aminopyridoxal-5’-phosphate); JZR (hexyl D-glucopyranoside); OLA (oleic acid); YOF (3-fluorotyrosine); OM2 ((R)-[2-[[(2S)-butan-2-yl]carbamoyl]phenyl]methyl-[(4-carboxy-1,3-benzodioxol-5-yl)methyl]-methyl-azanium); and OZR (boc-ps0-phe-gln-phe-NH2).

**Figure 3.**
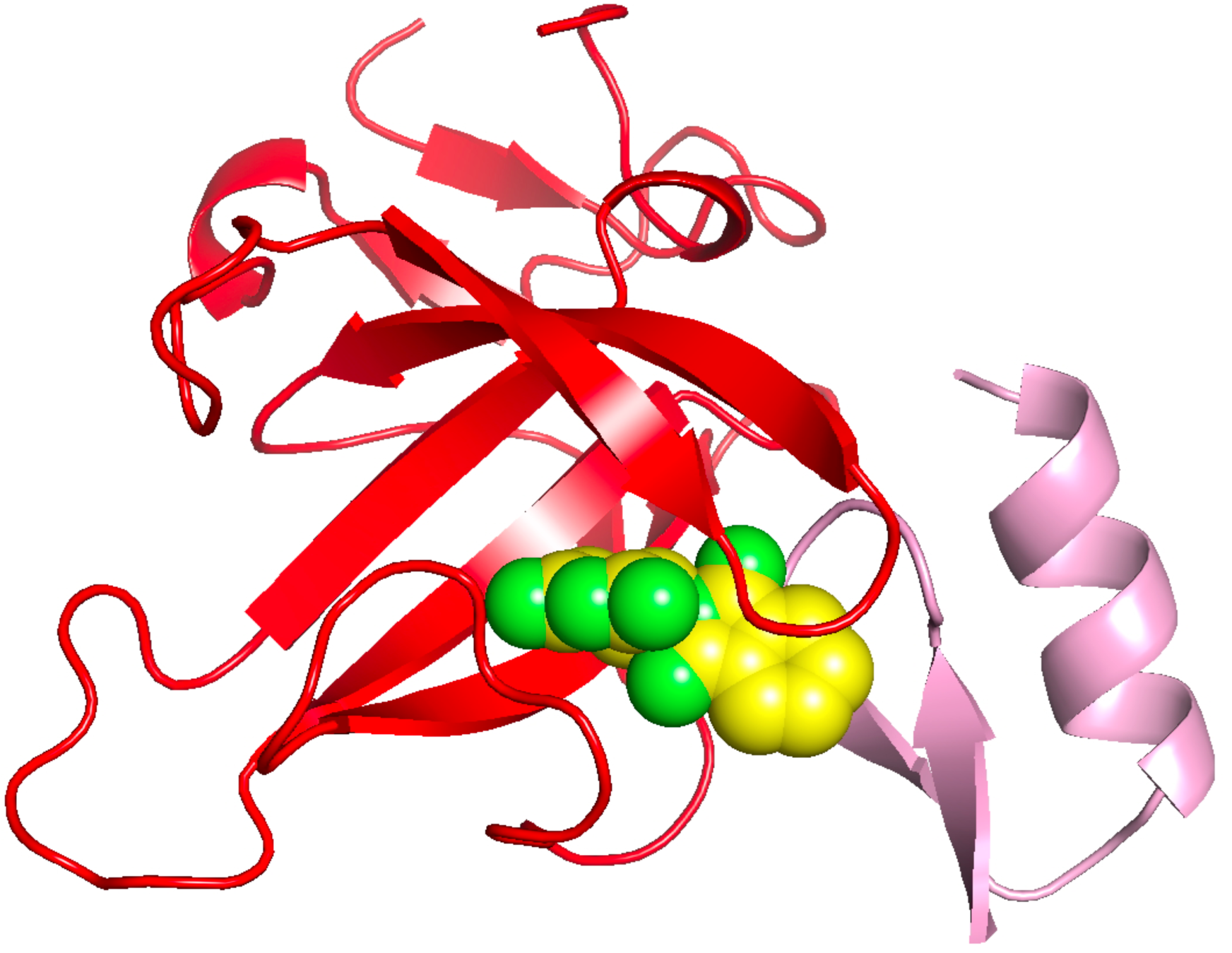
Predicted binding pose of thalidomide (green) compared to the experimental pose in the interface adjacent region between Cereblon (red) and SALL4 in 7bqu (23).

Despite their potential, rational discovery and design of molecular glues remain an unsolved problem, largely due to the complexity of PPI surfaces, the dynamic nature of ternary complex formation, the absence of predictive computational or statistical frameworks and the traditional emphasis on two E3 ligases, i.e. VHL and Cereblon (6). Current approaches heavily rely on trial-and-error or serendipitous screening, which is both inefficient and difficult to scale.

A robust, unbiased computational platform capable of systematically integrating structural, biophysical, and chemical data to model and predict molecular glue interactions would represent a transformative step forward. Such a platform could systematically identify new glueable targets, prioritize compound libraries, and optimize lead compounds with greater precision, thus accelerating the development of molecular glues across therapeutic areas.

Our work establishes such an unbiased computation guided approach for molecular glue discovery, decoupling design from known degrader scaffolds and specific ligase dependencies. By expanding the accessible E3 ligase landscape beyond CRBN and VHL and enabling rational targeting of interface adjacent binding pockets for an arbitrary collection of possible protein partners, our platform significantly broadens the scope and precision of targeted protein degradation, thus offering a transformative step toward systematic and mechanism-driven molecular glue development for diverse therapeutic applications.

## Results

### Examples of molecular glues in the PDB

As a first step, we analyzed interface-adjacent molecules in native protein dimers deposited in the Protein Data Bank (PDB). While not all of these ligands are molecular glues in the strict sense—that is, they do not induce novel interactions between proteins that otherwise would not associate—they nonetheless play an analogous physiological role by stabilizing protein–protein interactions. Their co-crystallization at interface-adjacent pockets (IAPs) indicates that their binding free energy is favorable. In SI, LIST.glues_pdb.txt, we provide the complete catalogue of such non-metal ligands found in the PDB, with methodological details described in Materials and Methods. In total, 1665 ligands were identified as IAP binding small-molecules with potential to act as molecular glues. The most common is glycerol, followed by ethyl 4-azanyl-3 bromanyl-benzoate. Another interesting molecule is pyroxidal-5’-phosphate, the active form of vitamin B6 (17). Small organic molecules such as acetate and citric acid are observed at interfaces, along with a diverse collection of phospholipids that preferentially localize adjacent to protein dimeric interfaces. In addition, we found that six metal ions, Cd, Cu, Fe, Mg, Ni and Zn are IAP binding ligands. Taken together, these findings highlight that interface-adjacent pocket (AIP) binding small molecules represent a mechanistically diverse group of ligands that can potentially help stabilize multimeric protein assemblies and thus act as molecular glues. Furthermore, this analysis highlights the potential of these interface adjacent molecules to serve as a library of fragments or potential cores that would form the starting point for development of molecular glues by bioisosteric replacements and scaffold-hopping among the protein pairs. These changes would potentially serve to introduce favorable changes in molecular shape and size, electronic properties, pKa, and polarizability that will potentially facilitate tighter interactions across the protein pairs. Ts, they could transform transient interactions into permanent ones with reduced *k_off_* for the bimolecular complexes.

A natural question is: what fraction of all PDB ligands function in this capacity? We have found that IAP binding ligands account for roughly 12% of all bound ligands in solved protein dimers. They span a wide spectrum, ranging from large cofactors such as hemes, to less physiologically relevant entities such as sulfate ions that may or may not mediate meaningful interactions. Notably, some metal ions—including zinc, iron, and cadmium—reside at protein interfaces, clearly stabilizing complexes and often contributing to enzymatic function. This information on interfacial metal ions could be leveraged to elucidate the principles of molecular glue design. Protein-metal ion binding affinity is dictated by Irving-Williams (IW) series (Mn^2+^<Fe^2+^<Co^2+^<Ni^2+^<Cu^2+^>Zn^2+^) with most metalloenzymes complexing with Zn^2+^ on one end of the spectrum and most metal-dependent enzymes binding to Mn^2+^/Mg^2+^ on the other end of the spectrum (18). The aspect of cognate metal speciation is a function of protein dynamics, non-equilibrium kinetic barriers and natural selection rules that dictate that the tightest binding metals are those that are least available (19). In fact, this know-how has been leveraged to design appropriate metal ion selectivity in protein design (7). A systematic analysis of interface adjacent metal ions suggests the following: From most to least frequent are Fe(10)>Cd(9)>Cu(7) and Mg(7)>Ni(4)>Zn(3), where the parenthesis indicates the number of examples found in the PDB. We note that while there are too few examples to tease apart the ground rules for interfacial metal ion recognition, they are roughly in the same inverse order as the IW series.

There are a total of 850 clusters when the ligands are clustered with a Tanimoto coefficient threshold of 0.8 using Daylight fingerprints (20). 19 dominant clusters that contain 10 or more members. The cluster centroids for the top ten clusters along with the number of members per cluster are shown in Figure 2B. The ligand clusters encompass a chemically diverse set of entities, ranging from highly charged groups to neutral scaffolds. A prominent feature is the prevalence of oxygen-rich functionalities which contribute polarity and hydrogen-bonding potential. Excluding the polyethylene glycols—the fourth largest cluster—many display drug-like characteristics. In addition, a significant subset is lipid-like with long hydrocarbon tails. This distribution is consistent with observations from the Protein Data Bank, where hydrocarbons frequently occupy interface-adjacent pockets as well as within protein– protein interfaces, exploiting their conformational flexibility to adapt to local stereochemistry (21). It should be emphasized that a substantial fraction of these ligand clusters represent monovalent small molecules with molecular weights of less than 500 Da, characteristic features of a molecular glue. Prominent examples include fluorotyrosine-like, amino pyridoxal phosphate-like and hexyl beta-D-glucopyranoside-like molecules. The complete ligand set is provided in the SI LIST.glues_clustered_0.8.txt.

### Validation of GlueFinder on the native protein dimers

As a preliminary step toward validating the utility of GlueFinder for predicting ligands that bind to pockets adjacent to protein–protein interfaces, we constructed a benchmark set of 238 solved human protein dimer structures from the PDB (see Materials and Methods for details). Since these dimers already have experimentally determined structures, this test evaluates the algorithm’s ability to predict and quantify GlueFinder’s precision and recall and binding pose prediction quality. As summarized in Table 1, we considered two scenarios. In the first, cases were restricted to those where APoc (22) assigned p-value of the template ligand–target pocket structural similarity score was less than 0.05. Within this set, the most stringent condition required template ligands to reside in proteins with no more than 30% sequence identity to their template protein with either monomer of the target dimer. Under these conditions, the recall was ∼24% with an average center-of-mass root-mean-square-deviation, RMSD, of 3.01 Å. Relaxing the sequence identity threshold to 40%—beyond which results were largely insensitive—increased coverage to slightly under one-third of the database, with an average RMSD of 2.82 Å. In the second analysis, we applied a more stringent criterion of pocket similarity with a p-value :< 0.01, again restricting template ligands to ≤ 30% sequence identity. Here, ∼7.4% of ligands were correctly matched, with an average RMSD of 2.57 Å. Increasing the sequence identity threshold to 40% yielded an average RMSD ≤ 2.39 Å, and the coverage increased to 12%. Although the overall coverage was lower than in the more permissive case (p-value < 0.05), the increased accuracy was modest, suggesting that an APoc p-value cutoff of 0.05 provides a robust balance of coverage and precision. These findings establish a practical framework for employing GlueFinder in virtual ligand screening, which we will next expand upon.

**Table 1.** Analysis of the ability to predict native glue-like ligands in the 238 protein dimer benchmark set.

| **Template**<br>**sequence**<br>**identity cutoff** | **Fraction of**<br>**native ligands**<br>**predicted** | **Average**<br>**predicted**<br>**precision per**<br>**native ligand** | **Average RMSD of**<br>**the predicted native**<br>**ligand's center of**<br>**mass (Å)** |
| --- | --- | --- | --- |
| **p-value $\leq 0.05$** |  |  |  |
| 30% | 0.24 (58/238) | 0.16 | 3.01 |
| 40% | 0.32(77/238) | 0.19 | 2.82 |
| 50% | 0.32(77/238) | 0.19 | 2.81 |
| 70% | 0.33 (79/238) | 0.19 | 2.74 |
| **p-value $\leq 0.01$** |  |  |  |
| 30% | 0.074 (20/238) | 0.27 | 2.57 |
| 40% | 0.12(41/238) | 0.28 | 2.39 |
| 50% | 0.17(41/238) | 0.26 | 2.39 |
| 70% | 0.18(42/238) | 0.28 | 2.34 |

By way of illustration of the accuracy of the approach, as shown in Figure 3, we successfully predicted that thalidomide (EF2) binds to the complex of CRBN to SALL4 (23), in agreement with experiment (PDB ID: 7BQU). The pose of EF2 is remarkably well predicted. This is an interesting case as the thalidomide pose is provided by the 5amk which is the monomeric structure of Cereblon. Indeed, one mechanism of ligand selection is to look for monomeric proteins that have a bound ligand which is partly exposed to the solvent. The solvent exposed piece can then interact with the appropriate pocket to generate binding of the two protein partners.

### Application to human dimers without known bound interface adjacent ligands

We next applied GlueFinder to a benchmark set of 2,682 human protein dimers from the PDB to evaluate its ability to predict intermolecular glues for a diverse collection of human proteins, the “Dimer” set. The goals of this analysis were twofold: (i) to determine the fraction of dimers that could be predicted to contain at least one ligand that putatively binds adjacent to the protein–protein interface, and (ii) to assess the predictive performance of the method in terms of precision and recall. As shown in Table 2, GlueFinder identified IAP ligands for 78% of native dimers, indicating that intermolecular glue candidates are potentially widespread across human protein complexes. In the absence of a target-template sequence identity cutoff, at a predicted binding precision threshold of 0.15, 2,089 proteins have a predicted binding precision of 0.18, with the best average binding precision of 0.24. The black curve in Figure 4 shows the cumulative histogram of the predicted binding precision. At a template sequence identity cutoff of 30%, and a minimum predicted precision of 0.05, 2,083 dimers are predicted to have interface adjacent binding ligands (see SI, dimers_0.30/stat_analyze.dimer_0.15_0.30). The average binding precision and fraction of known interface adjacent binding ligands is virtually the same as when a permissive sequence identity cutoff is used. The red curve in Figure 4 shows the plot of the cumulative fraction of predicted ligands versus the predicted binding precision. While the average precision is 18%, roughly 13% of the ligands have a predicted binding precision above 20%. This result suggests that a substantial fraction are likely to represent true positives, with the best having a predicted binding precision of 0.59, thereby supporting the utility of GlueFinder in prioritizing small molecules for subsequent experimental validation of their glueing potential. Interestingly, the cumulative fraction of cases with a predicted binding precision is very insensitive to template sequence identity cutoff. Indeed, anticipating the results below, the cumulative fraction of cases versus predicted precision curves are very similar for all considered scenarios.

**Figure 4.**
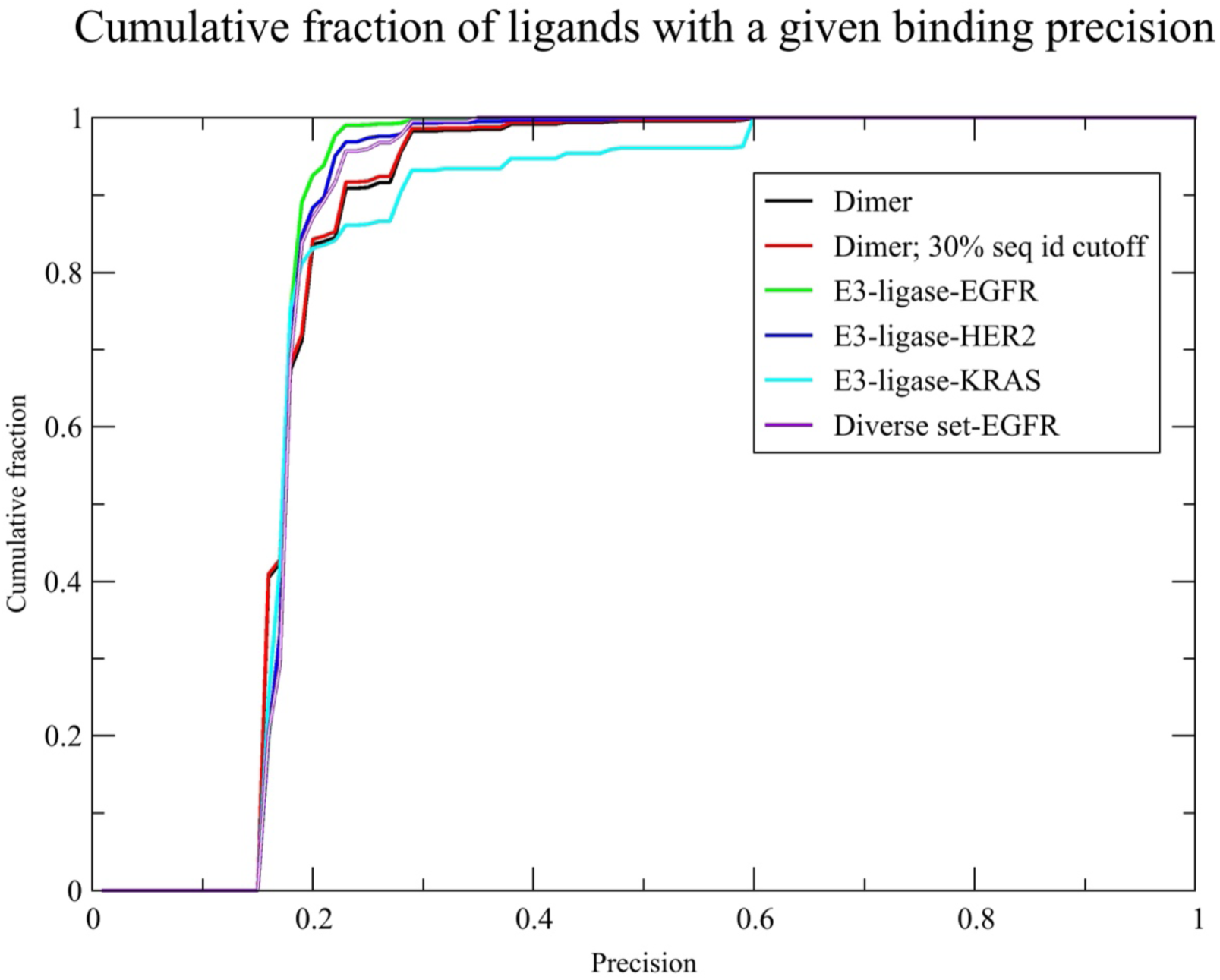
Plot of the cumulative binding precision for the six cases considered. The black line shows the set of native templates without known interfacial adjacent ligands and no template sequence identity cutoff period. The redline shows the set of native dimers with a 30% template sequence cutoff. The green line is the E3-ligase-EGFR l results. The blue line is the E3-ligase-HER2 results, the teal line is the E3-ligase-ligase results. The purple line is the diverse set-EGFR results.

**Table 2.** Average recall and precision for the various sets of protein dimers.

| **Template sequence identity cutoff used** | **Fraction of proteins for which glue molecules are predicted** | **Average precision of predicted ligand** | **Average fraction of interface adjacent native ligands** | **Average of best predicted binding precision (maximum value)** |
| --- | --- | --- | --- | --- |
| **Native human protein dimers** |  |  |  |  |
| No template cutoff | 0.78 (2101/2682) | 18% | 0.76 | 0.24 (0.59) |
| 30% template sequence identity cutoff | 0.78 (2083) | 18% | 0.76 | 0.29 (0.59) |
| **E3-ligases with EGFR** |  |  |  |  |
| No template cutoff | 0.83 (24/29) | 0.18 | 0.69 | 0.25 (0.38) |
| **E3-ligases with HER2** |  |  |  |  |
| No template cutoff | 0.98 (111/113) | 0.18 | 0.64 | 0.24 (0.59) |
| **E3-ligases with KRAS** |  |  |  |  |
| No template cutoff | 0.48 (148/31) | 0.20 | 0.80 | 0.21 (0.59<sub>xw</sub>) |
| **Diverse proteins with EGFR** |  |  |  |  |
| No template cutoff | 0.36(13/36) | 0.18 | 0.83 | 0.23 (0.35) |

<sup>a</sup> Predicted binding precision threshold of 0.15

Next, we considered a subset of the 10 most frequently predicted ligands obtained without any restrictions on the relationship of the target and template proteins; all are known interface adjacent binding ligands. Interestingly, the same top ligands are predicted even when the closest target-template sequence identity is :< 30%. Some of the top ligands are provided in Table 3, column 1. We present some of these ligands in detail as they will often be found in subsequently analyzed predictive situations involving E3-ligases as well as other classes of molecules. Thus, it is important to establish the plausibility of these results.

**Table 3.** Subset of the frequently selected putative molecular glues for E3-ligase with EGFR, HER2 and KRAS.

| Native dimers | EGFR | HER2 | KRAS |
| --- | --- | --- | --- |
| Name of ligand<br>(Number of dimers bound) |  |  |  |
| HEC<br>(Heme)<br>(1115)* | OLC<br>(oleic acid)<br>(19)* | MPD2<br>(methyl-2-4 pentane<br>diol)<br>(69)* | MLY<br>(N-methyl lysine)<br>(89)* |
| MPD<br>(2-methyl-2,4-pentanediol)<br>(919)* | LFA<br>(eicosane)<br>(18)* | CYC<br>(phycocyanobilin)<br>(43) | BCL<br>(bacteriochlorophyll)<br>(26)* |
| OLC<br>(oleic acid)<br>(627)* | BCL<br>(bacteriochlorophyll)<br>(18) | DMU(decyl-beta-d-<br>maltopyranoside)<br>(37) | MES<br>2-(n-morpholino)-<br>ethanesulfonic acid<br>(18)* |
| BCR<br>(beta-carotene)<br>(222) | SQD (1,2-di-o-acyl-3-o-<br>[6-deoxy-6-sulfo-alpha-d-glucopyranosyl]-sn-glycerol)(18)* | BCL<br>(bacteriochlorophyll)<br>(37) | DMU<br>decyl-beta-d-<br>maltopyranoside<br>(17) |
| PLM<br>(Palmitic acid)<br>(667)* | BCR(beta-carotene)(14)L2P | LMG<br>(1,2-distearoyl-<br>monogalactosyl-<br>diglyceride)(15) | LDA<br>lauryl dimethylamine-<br>n-oxide<br>(17)* |
| CIT<br>(Citric acid)<br>(526)* | L2P<br>(2,3-di-phytanyl-<br>glycerol)<br>(12) | B3K<br>(3s)-3,7-diamino-<br>heptanoic acid<br>(13) | SQD<br>sulfoquinovosyldiacyl<br>glycerol<br>(11) |
| CLR<br>(cholesterol)<br>(452)* | PEK<br>((1s)-2-([(2-aminoethoxy)(hydroxy)p<br>hosphoryl]oxy}-1-<br>[(stearoyloxy)methyl]eth<br>yl (5e,8e,11e,14e)-icosa-5,8,11,14-tetraenoate)<br>(9) | BLA<br>biliverdin ix alpha<br>(12) | LMG<br>1,2-distearoyl-<br>monogalactosyl-<br>diglyceride<br>(11)* |
| CYC<br>(phycocyanobilin)<br>(431) | PLM (palmitic acid)<br>(9)* | FTR<br>Flurotryptophane<br>(10) | DGD<br>digalactosyl diacyl<br>glycerol<br>(10)* |
| CDL<br>(cardiolipin)<br>(289)* | TGL<br>(tristearoylglycerol)<br>(7) | 78N<br>((2r)-2,3-dihydroxypropyl(7z)-<br>pentadec-7-enoate<br>(8) | TLA<br>Tartaric acid<br>(5)* |
| CLR<br>(cholesterol)<br>(15)* | PGV<br>(1R-2-([(2s)-2,3-dihydroxypropyl]oxy}{hy<br>droxy}phosphoryl]oxy}-<br>1-<br>[(palmitoyloxy)methyl]et<br>hyl (11e)-octadec-11-enoate)<br>(6) | XCP<br>(1S,2S)-2-aminocyclopentanecarb<br>oxylic acid<br>(8) | BNG<br>beta-D-glucopyranoside<br>(5)* |

\*Known interface adjacent binding ligand.

Among these is the heme group. Hemes cause ligand-induced receptor dimerization through irreversible bis-histidine ligation. Free heme binds to the TLR4/MD-2 complex and promotes TLR4 dimerization and signaling (24, 25). This is a glue-like effect because the ligand stabilizes a receptor-dimer interface. Thus, there is credible evidence that heme can act *glue-like* under certain circumstances, stabilizing or inducing multimeric assemblies (e.g., in TLR4 dimers).

Another interesting case is MPD (4s-2-methyl-2,4-pentanediol). Solved protein structures provide evidence that MPD is “glue-like,” i.e., it can sit adjacent to protein–protein interfaces or even promote oligomerization (26). In Ca²⁺-calmodulin, an MPD molecule mediates a lattice contact by bridging helices of neighboring molecules. MPD alone (10–30% v/v) can induce heptamer formation of *Staphylococcus aureus* α-hemolysin *in vitro*; two MPD molecules bind at sites that facilitate inter-protomer interactions and stabilize the pore. Thus, MPD promotes protein–protein assembly under high-concentration conditions (27). Furthermore, in an antibody–fluorescein Fab structure crystallized with MPD, a bound MPD molecule is trapped at the inter-domain interface just below the antigen site, consistent with a small-molecule occupying and effectively “gluing” an interface in its crystal structure (28).

Evidence that oleic acid might also act as a glue is found in the interface regions of a number of solved protein structures (21). Another long chain fatty acid is PLM, palmitic acid. Tri-palmitoylated lipopeptides glue TLR1–TLR2 (29). The immune agonist Pam3CSK4 contains three palmitoyl chains that occupy hydrophobic pockets in both TLR2 and TLR1 which physically bridge the heterodimer. This is an archetypal “glue-like” mechanism (29). A C16 fatty acyl chain can bridge Frizzled receptors upon Wnt binding. Structures of Frizzled cysteine-rich domains (CRDs) bound to palmitoleic acid show that the acyl chain spans two CRD monomers, implying Wnt’s lipid *glues* a Frizzled dimer (30). Another study concluded that palmitic acid directly binds MD-2, promotes MD-2/TLR4 complex formation, and possibly triggers downstream signaling (31, 32). Finally, there is a related precedent for lipid chains gluing TLR4 dimers. The TLR4–MD-2–LPS co-crystal structure shows lipid A’s acyl chains filling the MD-2 pocket and bridging two TLR4–MD-2 complexes into the signaling-competent “M-shaped” dimer. Once again, a lipid-mediated “glue” is employed (33).

In the case of citric acid and its close derivatives, citrate is a classic allosteric activator of acetyl-CoA carboxylase (ACC1/ACC2) that induces dimerization/polymerization into active filaments (34). Furthermore, for β-carotene, there is direct structural evidence that it behaves as a molecular glue within native photosynthetic protein complexes, where it helps secure contacts between membrane-protein subunits (35). β-carotene (and other carotenoids) is a structural cofactor embedded in the membrane that stabilizes protein–protein contacts; that is, it acts very “glue-like,” in a biophysical sense (36).

As another example, there is strong structural and functional evidence that cardiolipin (CDL) molecules occupy interfaces and stabilize assemblies, notably respiratory-chain supercomplexes and ATP synthase dimer; thus, they act as bona-fide lipid glues in the inner mitochondrial membrane (37). Cardiolipin binds in an interface adjacent pocket in 8ps0, an endosomal sodium/proton exchanger whose dimeric state is dictated by the presence of cardiolipin (38). Below in Figure 5C, we show a predicted example of how CDL could stablize the solute carrier 13 dimer (39). Finally, we note that in membranes, cholesterol can behave as a molecular glue. Multiple structural and mechanistic studies show cholesterol molecules sitting at protein–protein interfaces and stabilizing dimers/arrays, i.e., functioning as interfacial cofactors rather than merely altering bulk membrane properties (40). However, evidence for cholesterol gluing soluble proteins in the cytosol is lacking to date.

**Figure 5.**
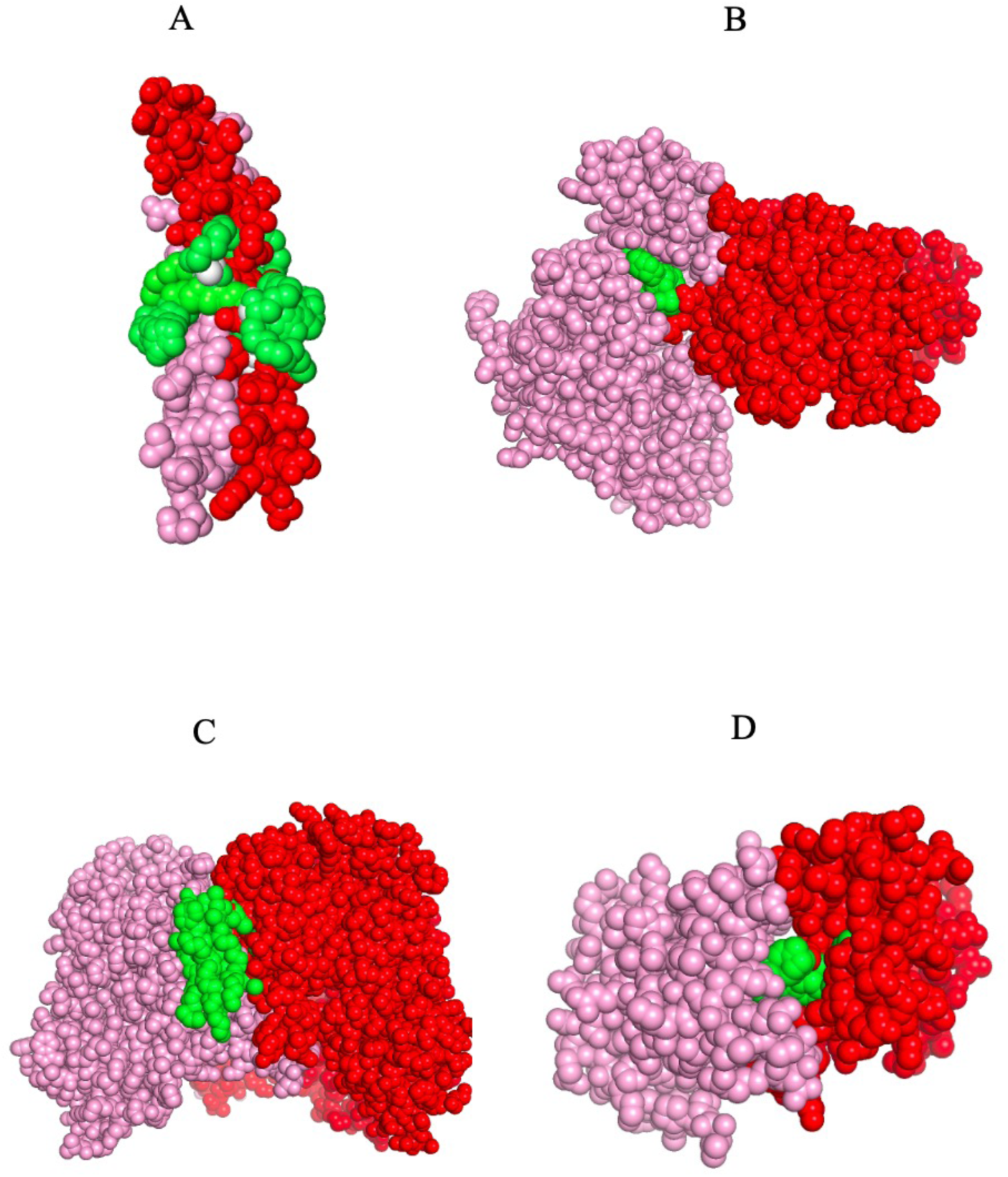
Binding poses of the ensemble of ligand conformations predicted to bind to the indicated protein dimer chain A is in pink, chain B in red and the dimer and the binding ligand is in green. A. Human striatin-3 coiled coil domain, 4n6j (41) binding TGL (tristearoylglycerol). B. ERK3, 6yll (42, 43), binding to STU (staurosporine). C. Solute carrier 13, 8w6h, (39) binding to CDL (cardiolipin). D. Fifth bromodomain of poly-bromodomain containing protein 1, 6g0j, (11) binding to R78 (4-([(7R)-8-cyclopentyl-7-ethyl-5-methyl-6-oxo-5,6,7,8-tetrahydropteridin-2-yl]amino}-3-methoxy-N-(1-methylpiperidin-4-yl)benzamide.

Figure 5 shows high-confidence representative examples of binding poses of small molecule ligands/ that are predicted to bind in the protein interface adjacent region. Figure 5A considers the human striatin 3-coil-coil domain, PDB id 4n6j (41), that is predicted to bind to tristeroglycerol (TGL). What is particularly interesting is that tristeroglycerol is a long three branched star-like molecule that literally coils around the coil-coil structure of 6n6j much like a snake constricts its prey. This suggests a possible new type of molecular glue constrictor. Figure 5B shows the binding of ERK1, whose PDB code is 6yll (42, 43), that is predicted to bind to staurosporine (STU). Located in an IAP, it putatively acts to help facilitate the intermolecular interaction. Figure 5C shows the predicted binding of cardiolipin to solute-character protein 8w6h (39). Cardiolipin is a long-chain molecule that contains four branches, and contrary to what one might imagine, it folds back upon itself to fill the crevice between the interface of two proteins. Figure 5D shows the fifth bromodomain of poly-bromodomain containing protein 1, 6g0j (11), which is predicted to bind to R78, a benzamide. Across cases, the predicted conformations are sterically and chemically reasonable, with ligands complementing local shape and polarity to buttress protein–protein contacts in the face-adjacent region.

### Putative glues between E3-ligases EGFR, HER2 and KRAS

In this section, we restrict ourselves to the case where AF3Complex (44) generates reasonably confident structure predictions (IS-score>0.30). As a consequence GlueFinder’s maximal coverage is limited by the ability of AF2Complex to generate reasonably confident quaternary structure predictions. In the next section, we then explore what happens if we use AF3Complex to generate possible quaternary structures, even if they are unlikely to occur in the absence of a molecular glue. This is of course the domain where molecular glues are needed.

For the case of binding EGFR to ligases, AF2Complex was able to provide 29 predicted dimers (whose combined length is <1700 residues) that contain at least five residues on each chain that are involved in the protein-protein interface. Thus, GlueFinder’s maximal coverage limits the ability of AF2Complex to generate reasonably confident quaternary structure predictions. 24/29 of these complexes have at least one predicted molecular glue whose binding precision is ≥ 0.15 (see SI/EGFR/LIST.targets_0.15.EGFR.txt). Furthermore, the average best binding precision is 0.24, with the best predicted ligand having a binding a precision of 0.59. The cumulative fraction of ligands with a given binding precision is shown in the green curve of Figure 4. The histogram of the set of binding ligands per protein is provided in SI, LIGANDS.EGFR.txt. There is a total of 167 different types of ligands bound, of which 115 (69%) are known interface adjacent binding ligands indicated by “**”. The corresponding full set of binding ligands and their predicted precision per protein in EGFR/summary/XX.summary with X being the name of the protein dimer. The phylogenetic tree of the set of 24 E3 ligases along with their relationship to canonical molecular glues is shown in Figure 6.

**Figure 6.**
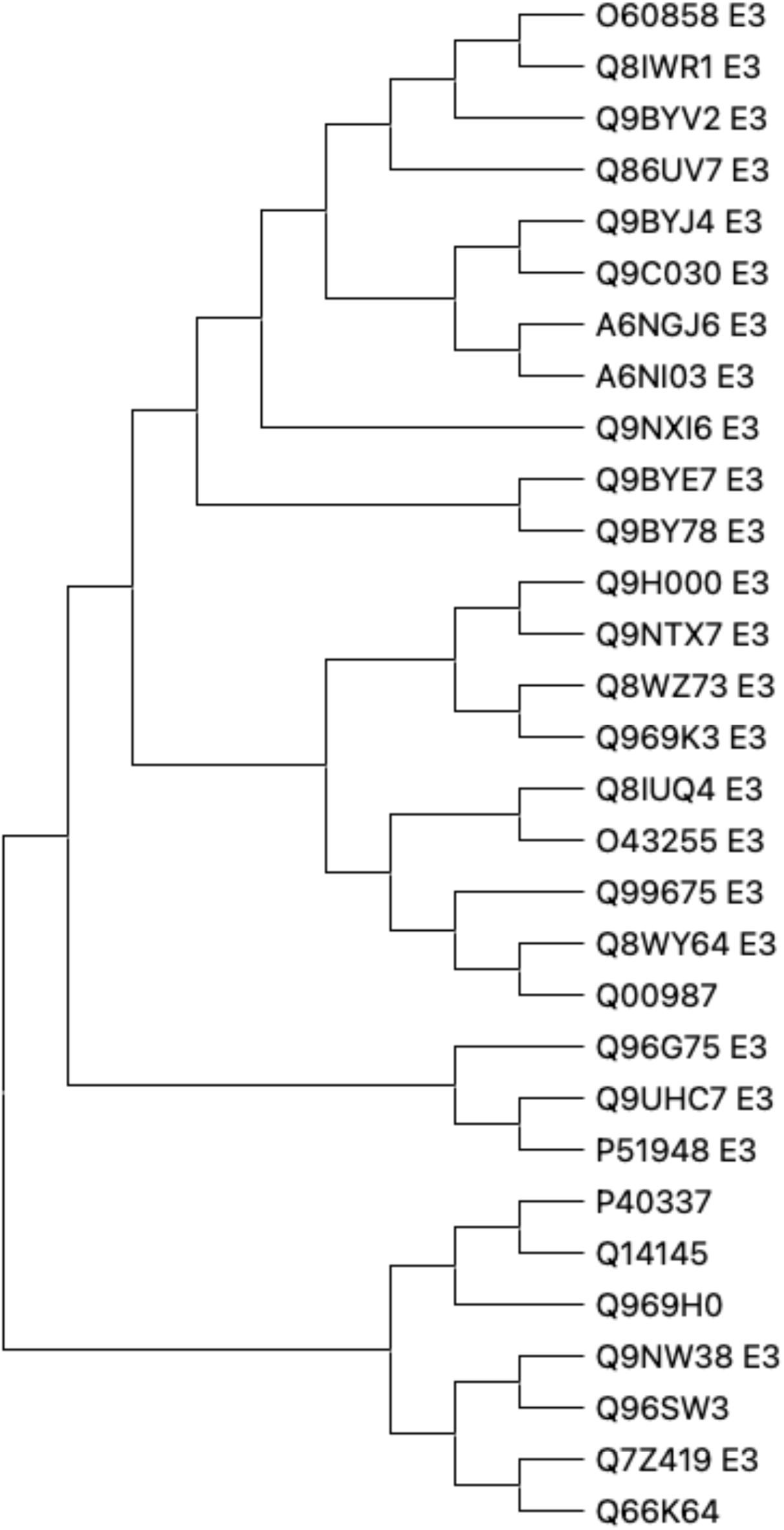
Phylogenetic tree of the 24 E3-ligases predicted to interact with EGFR assisted by a molecular glue-like metabolite are shown.

For E3-ligase-EGFR dimers, a subset of interesting putative molecular glues is presented in column 2 of Table 3. The sulfolipid SQG, (sulfoquinovosyldiacylglycerol) can play a bona fide **“**lipid glue” role by bridging protein subunits and stabilizing/determining dimerization in photosynthetic complexes (45). High-resolution structures reveal two SQG molecules (SQD-1 and SQD-2) at the monomer–monomer interface. Each forms specific hydrogen bonds to residues on different subunits across the two monomers (e.g., SQD-1 links D1 of one monomer to CP47 of the other). Removing SQD breaks these bridges and shifts PSII toward monomers, i.e., it destabilizes the dimer. This is exactly a glue-like function at the protein–protein interface (45). Decades of PSII work conclude that bound lipids mediate protein–protein and monomer–monomer contacts within these complexes (46). SQD is one of the lipids repeatedly found at such oligomerization interfaces. Finally, other thylakoid lipids sometimes *are* explicitly described as a “molecular glue” (e.g., phosphatidylglycerol is proposed to glue LHCII monomers), showing that structural lipids can function as glues in native membrane assemblies (47).

In the case of HER2, as shown in columns 2 and 3 of Table 3, GlueFinder was able to predict binding ligands for 111/113 dimers (see SI/HER2/LIST.targets_0.15.HER2.txt) with a predicted binding precision at or above 0.15. The cumulative fraction of ligands with a given predicted binding precision is shown in the blue curve of Figure 4. Furthermore, the histogram of the set of binding ligands per protein is provided in SI, HER2/LIGANDS.HER2.txt, and the corresponding full set of binding ligands and their predicted precision per protein is in HER2/summary/XX.summary with “XX” being the name of the protein dimer. A total of 237 different types of putative molecular glues are predicted with ∼64% (151/237) being known interface adjacent ligands. The phylogenetic tree of the 113 E3 ligases having predicted glues for HER2 along with their relationship to canonical molecular glues is shown in SI, Figure S3. The most frequent novel putative glue is phycocyanobilin that binds to 43/111 E3-ligase-HER2 complexes. In addition, MPD (2 methyl-2-4 pentane diol), a known interface adjacent binding ligand, is among the most frequent putative glues. MPD induces assembly of α-hemolysin into a heptamer (27). A study that crystallized *Staphylococcus aureus* α-hemolysin from monomeric proteins found that MPD alone (no membranes/detergent) drove formation of the SDS-resistant heptameric pore. There were two localized MPD molecules per protomer near Trp179; SDS-PAGE showed efficient oligomerization. MPD facilitates and stabilizes inter-protomer contacts by mimicking lipid head-group interactions and providing a membrane-like environment. Furthermore, MPD directly participates in a protein–protein interface associated with cholera toxin B assembly (48). In a designed CTB pentamer system, the crystal structure revealed that MPD sites are located in the interface between pentamers (48). Another novel ligand is LMG that can act as an IAP “glue” that stabilizes contacts between membrane-protein subunits in photosynthetic complexes. Multiple high-resolution structures and biophysical studies show LMG molecules wedged at subunit interfaces or mediating hydrogen-bond networks that hold complexes together (36). Furthermore, in at least one system, BLA, behaves “glue-like**”** (49). A cryo-EM structure of the human mitochondrial transporter ABCB10 shows BLA sitting in a pocket of one protomer and bridging the interface to the opposite protomer via hydrogen bonds, stabilizing a more closed, catalytically competent conformation. Thus, it is functionally acting as an interfacial cofactor that holds the dimer together (49). BLA binding also brings the two nucleotide-binding domains closer (from ∼30 Å to ∼13 Å), consistent with stabilization of the closed state.

In the case of E3 ligases putatively binding to KRAS, as shown in Table 3, 148/321 predicted types of complexes, have glues that are predicted to bind (see SI/KRAS/LIST.targets_0.15.txt) with a precision ≥ 0.15, with the best average precision of 0.21. Again, we only consider predictions where at least two examples are found. The full list of putative glues for each of the 321 screened E3-ligases (having at least one example) is found in SI/KRAS/summary/**.summary. The highest precision ligand has a value of 0.59. The cumulative fraction of ligands with a given binding precision is shown in the teal curve in Figure 4. Of the 127 putative glues, 93 are known interface adjacent binding ligands, with representative examples shown in column 4 of Table 3. DMU is a novel putative glue with some literature evidence that it can help keep specific membrane protein complexes intact by occupying native lipid sites in proteins involved in the photosystem II complex (50). The gluing potential of SQD has been mentioned above. DGD does not yet have any approved medical uses, but it does have anti-inflammatory activity in mouse models (51) and has anticancer activity/synergistic activity with doxorubicin in melanoma cells (52). The origin of this activity is unknown at the moment.

### Molecular glues for the Diverse-EGFR set

In a harder test set, in the Diverse-EGFR set, we predicted the quaternary structure of EGFR in complex with a collection of another 25 proteins, see Table 4. The cumulative precision of the predicted ligands is found in the purple curve of Figure 4. Here, we deliberately considered cases where ***these proteins are not predicted*** by AF2Complex to interact with EGFR in the absence of a ligand; rather, we are employing AF2Complex to predict interface adjacent pockets in structures which might occur with very low binding affinity. Table 4 provides an exceptionally diverse set of protein families predicted to associate with it through stabilization by intermolecular glues. These include homeobox and forkhead box transcription factors, aminotransferases, DNA and RNA polymerases, viral and cellular antigens, RAS-related GTPases, guanine-binding proteins, aquaporins, additional classes of transcription factors, membrane proteins, kallikreins, RNA-binding proteins, DNA–RNA-binding proteins, NF-κB–inhibitory RASD proteins, and zinc finger proteins. Collectively, this striking breadth of potential interactors suggests that EGFR can be modulated within a heterogeneous repertoire of protein contexts. Such diversity underscores the possibility of unanticipated modes of regulation and highlights opportunities for novel therapeutic interventions, which will be elaborated upon below.

**Table 4.**
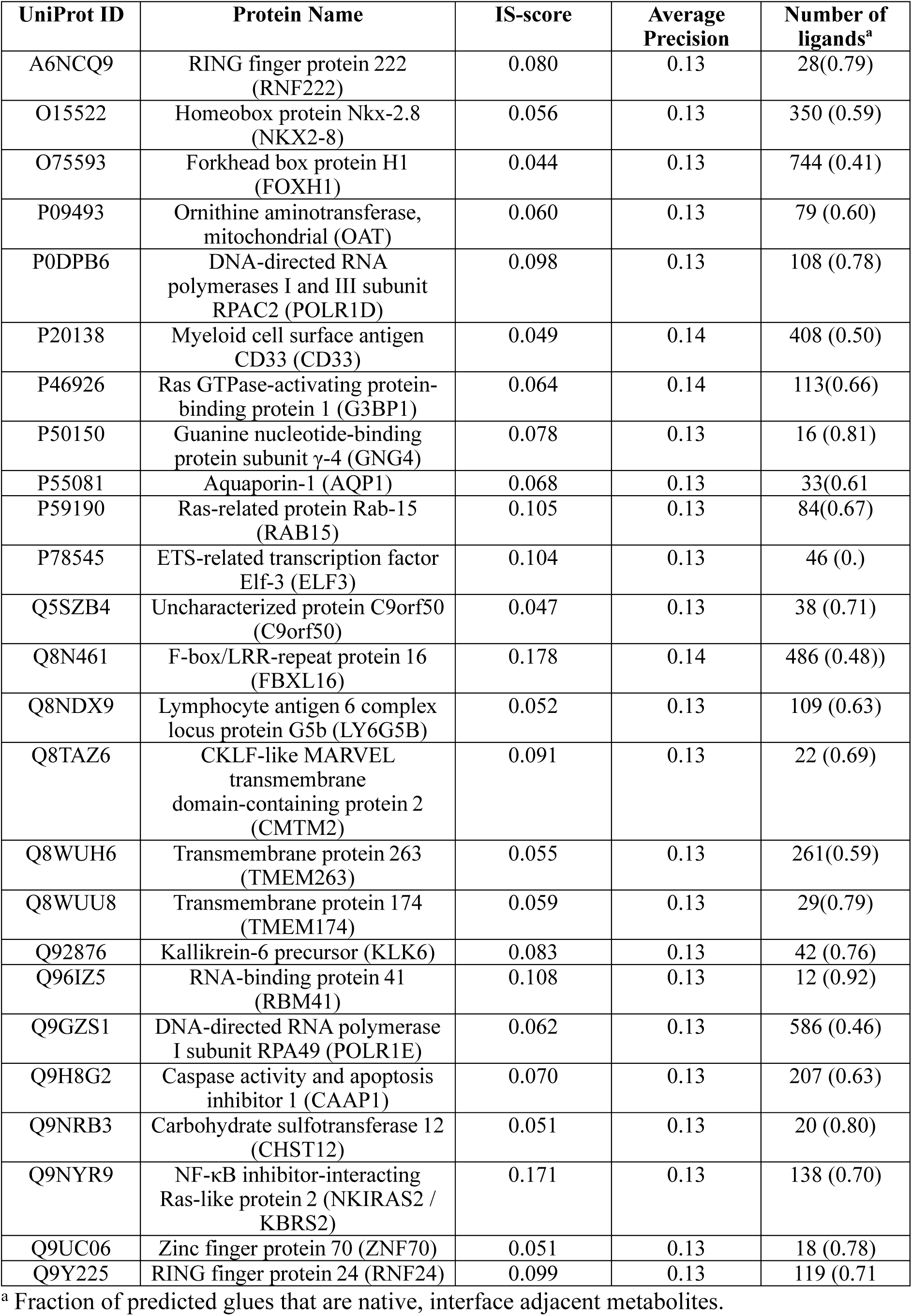
Results for the Diverse-EGFR set.

The total set of binding ligands per protein is provided in SI, diverse_EGFR/LIGANDS.diverse_EGFR.txt. The full set of ligands and the predicted precision per protein in *. summary. * is the name of the protein dimer along with the predicted binding glues and binding precision that are found in SI, diverse_EGFR/summary/XX.summary. As shown in Table 2, the average predicted precision is 0.18, with 83% of the predicted putative glues in the native interface adjacent ligand library. The average of the best predicted binding precision is 0.23, with the absolute best predicted binding precision of 0.35. As previously, ligands not in the native IAP ligand library include CYC (phycocyanobilin), BCL (bacteriochlorophyll), BCR (beta-carotene), GLY (glycine), UMP (5 deoxyuridine 5’monophosphate), SQD (1,2-di-o-acyl-3-o-[6-deoxy-6-sulfo-alpha-d-glucopyranosyl]-sn-glycerol) and AK6 (2-oxoglutaric acid).

### Examination of EGFR with canonical E3 ligases having known molecular glues

For EGFR, none of the four E3 ligases which have molecular glues to other targets, Cereblon (Q96SW2) (53), VHL (P40337) (54), MDM2 (Q00987) (55), DCAF15 (Q66K64) (56), are predicted to form stable dimers without a molecular glue by AF2Complex. For Cereblon, citric acid (SAH, adenosyl homocysteine) is predicted to be a molecular glue with EGFR with a precision of 0.21 (0.19). For VHL, NAG, 2-acetamido-2-deoxy-beta-D-glucopyranose (BCR, beta-carotene), is predicted to be a molecular glue with EGFR with a precision of 0.29 (0.17). For MDM2, Y01, cholesterol hemi succinate (OLC, 2R-2,3-dihydroxypropyl (9Z)-octadec-9-enoate) is predicted to act as a molecular glue with EGFR with a precision of 0.18 (0.17). For DCAF5, a top putative molecular glue is CIT (BTB, 2-[bis- (2-hydroxy-ethyl)-amino]-2-hydroxymethyl-propane-1,3-diol) with a predicted binding precision of 0.19 (0.14).

## Discussion

The development of molecular glue degraders opens a transformative avenue in targeted protein degradation, enabling modulation of previously “undruggable” proteins by promoting proximity-induced ubiquitination and subsequent proteasomal degradation. Despite their therapeutic promise, molecular glues have largely been discovered serendipitously, with rational design approaches still in their infancy. A major bottleneck is the field’s overreliance on a narrow set of E3 ligases, primarily Cereblon (CRBN) and von Hippel–Lindau (VHL), which limits the diversity of targetable substrates and creates resistance risks through ligase saturation or mutation. Expanding the set of proteins beyond these traditional ligases is essential for fully realizing the potential of molecular glues as a generalizable therapeutic modality.

To achieve this, there is a critical need for a rational, unbiased platform that can systematically explore the entire E3 ligase landscape and identify potential molecular glue opportunities across a broad range of protein targets. A key innovation in this direction is the global structural and statistical analysis of protein–protein interfaces and their adjacent ligand-binding events across the PDB. By identifying ligands that naturally localize near protein– protein interfaces—particularly those that bridge or stabilize two proteins—one can infer structural and chemical principles that favor ternary complex formation. These protein interface adjacent ligand-bound pockets often represent latent opportunities for molecular glue design, even if they were not originally identified in the context of degradation.

This ligand-centric, interface-aware strategy provides a scalable means to prioritize small molecules and protein pairs for experimental validation, offering a stark contrast to trial-and-error screening approaches. Importantly, such a method is agnostic to the identity of the E3 ligase, enabling exploration of the full human E3 ligase repertoire—over 600 ligases— rather than restricting design to a few well-characterized examples. This not only diversifies potential degradation mechanisms but also opens avenues for tissue-specific or disease-selective targeting by leveraging E3 ligase expression patterns.

Recent advances in structural biology, particularly AlphaFold 2 (57), AlphaFold-Multimer (58), AF2Complex (59), AlphaFold 3 (60) and AF3Complex (61) now enable high-confidence predictions of quaternary protein structures at scale, including E3-ligase complexes for which no experimental structure exists. Moreover, even when a given set of proteins are not predicted to form inherently *stable* dimers, these algorithms can generate reasonable structures to which a ligand could bind to its interface adjacent pocket, thereby stabilizing the complex. This after all is the goal of molecular glues. Integrating predicted POI–E3 complexes with statistical maps of interface-adjacent ligandability allows researchers to computationally screen for “druggable” interfacial pockets across theoretical ternary assemblies. This empowers a rational framework for glue design even when direct structural data is lacking, significantly expanding the scope of targetable interactions.

Our putative EGFR diverse protein interactors span transcriptional regulators (ETS-family factors and zinc-finger proteins), protein quality control proteins (RING-type E3 ligases), vesicle trafficking proteins (the RAS-related GTPase RAB5), and even DNA-associated enzymes (including polymerases). This diversity suggests that interface-adjacent small molecules (“molecular glues”) could be leveraged not only to stabilize cryptic EGFR complexes but also to reroute the information flow across levels of regulation—transcription, membrane trafficking, and proteostasis. For example, glues that couple ETS factors or other co-activators to EGFR-proximal chromatin machinery could upregulate EGFR expression under physiological, non-oncogenic contexts (e.g., tissue repair), whereas glues that recruit RING E3s to EGFR would favor ligase-directed ubiquitylation and turnover. Likewise, glues that bias EGFR’s endocytic itinerary via RAB5-proximal effectors could retune receptor surface dwell time and signaling duration. Conceptually, this extends synthetic-biology paradigms—where small molecules have been used to retask protein assemblies—to reengineer native mammalian complexes *in situ* for programmable gain- or loss-of-function.

We further note that the absence of supporting literature does not necessarily invalidate the predicted set of molecules as possible molecular glues. Rather, it may reflect their novelty and offers the potential to uncover previously unrecognized molecular glues. At the same time, one must acknowledge the possibility of over-prediction, wherein certain compounds bind so weakly that they fail to exert a meaningful structural or functional effect. In such cases, the ligand may not induce dimerization. Thus, while some predictions may represent bona fide novel glues, others may prove nonfunctional, underscoring the need for careful experimental validation.

Broadly speaking, two principal classes of molecular glues can be considered. The first comprises the so-called bridging compounds, here referred to as interface-adjacent pocket (IAP) ligands, which occupy the interfacial region between two interacting proteins and directly stabilize their association. The second class consists of allosteric glues, which act at a distance by inducing conformational rearrangements that expose previously buried hydrophobic surfaces, thereby promoting new protein–protein interactions. While the latter mechanism cannot be excluded, the current class of algorithms is specifically designed to identify the former type—that is, ligands residing in or near the protein–protein interface. Interestingly, even in systems such as CRBN, where one domain sometimes undergoes partial opening upon ligand binding, the ligand-binding domain itself remains largely structurally conserved, functioning effectively as a rigid body whose ligand pose is structurally conserved consistent with the assumptions underlying GlueFinder. Although the present framework does not explicitly predict domain cracking or other large-scale rearrangements—tasks more suited to structure prediction algorithms of the AlphaFold class—it remains applicable in cases where the rearranged domain still engages its partner through an interfacial or interface-adjacent ligand. Thus, while this approach does not yet provide a complete solution to the full spectrum of molecular glue mechanisms, it represents a significant step forward. Ongoing efforts aim to extend the method to predict allosteric conformational transitions, thereby addressing scenarios where ligand binding indirectly promotes or stabilizes protein–protein association.

Our work, by integrating global structural analysis with AI-powered structure prediction, provides a rational, predictive approach towards molecular glue discovery—one that does not depend on chance or historical bias. This paradigm shift holds the potential to unlock new degrader targets, harness underutilized E3 ligases, and create a scalable, mechanism-informed path for therapeutic development. It is through this convergence of computational insight and structural understanding that the full power of molecular glue degraders can be harnessed across the proteome.

## Materials and Methods

To demonstrate its generality, GlueFinder was applied under three scenarios with (22). A schematic overview of the methodology is shown in Figure 7. For cases with pre-existing experimental dimeric structures in the PDB, interface-adjacent pockets are identified and characterized using the Cavitator pocket alignment algorithm (22). For a pair of molecules whose quaternary structure is unknown, the structure of their complex is predicted using AF2Complex (AF2C) (59). Even when a confident protein–protein interaction cannot be predicted, we empirically observed that AF2C frequently generates reasonable interface-adjacent pocket poses suitable for small molecule screening. This scenario is particularly relevant for the design of molecular glues, where the interacting partners are unlikely to associate without a third molecule that stabilizes the complex and shifts the binding free energy toward the bound state. For such predicted structures, interface-adjacent pockets are first identified using Cavitator. Subsequently, the APoc protein pocket structure alignment algorithm (22) is applied to screen a library of small molecule pockets found in the entire Protein Data Bank (PDB). Candidate complexes are then ranked and subjected to virtual screening with the LIGMAP algorithm (62), which also predicts binding poses and provides associated binding precision estimates that the ligand will bind with at worst a 10 μM binding affinity.

**Figure 7.**
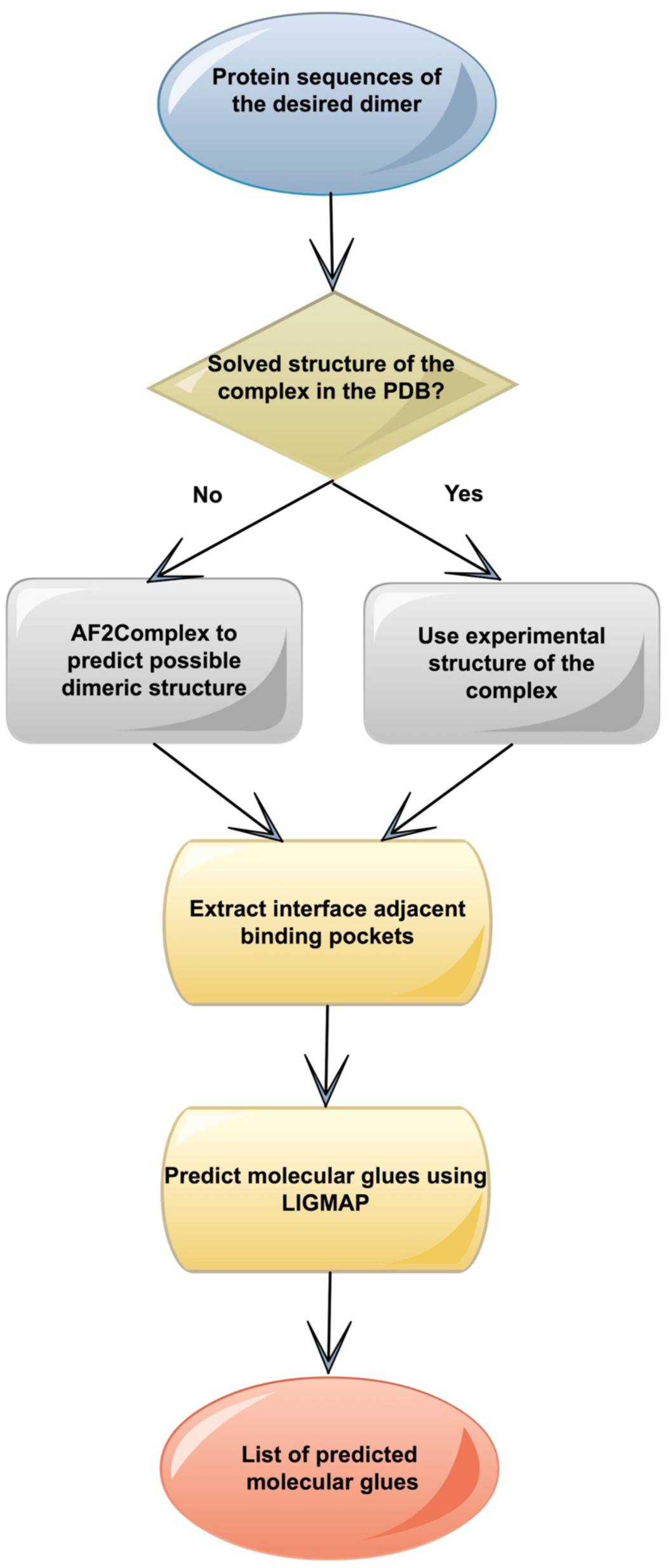
Overview of the GlueFinder molecular glue prediction algorithm.

### Databases

#### Native interface adjacent binding ligands

The entire Protein Data Bank (PDB) from September 2024 was scanned to identify dimeric complexes, irrespective of whether they were homodimers or heterodimers. For the purposes of this analysis, only dimers were considered, yielding a total of 63,091 cases (see SI LIST.fullpdb_chains_mult.dimers). To ensure that the interactions between chains represented genuine dimeric contacts, we required that each chain contribute at least five interacting residues. This criterion was chosen to provide a minimal likelihood that the observed interfaces correspond to physiologically meaningful interactions. From this curated set of dimers, we extracted all bound ligands and then restricted the analysis to those ligands that contacted at least five residues on each chain of the dimer. Application of these filters resulted in a set of 1,665 ligands, which are provided in the SI, LIG.glues.unique.txt; the dimeric proteins that contributed to this list are found in SI, LIST.fullpdb_chains_mult.dimers_hetcon_noMSE.sort.txt.

#### Library of solved human protein structures

All dimeric, human protein complexes in the Protein Data Bank (PDB) were extracted, yielding 15,680 structures. Of these, 238 are protein complexes with known interface adjacent ligands (see SI, LIST.benchmark_with_ligands.txt). This “native benchmark” set will be used to establish the precision and recall of GlueFinder as well as its ability to correctly position ligands in the interface adjacent pocket. To reduce redundancy and generate representative datasets, the full set of human PDB dimers were clustered at 70% sequence identity, yielding 2,695 complexes; see SI, LIST.clustered_70. Of these 2,563 have ligand binding predictions at a template sequence identity cutoff of 30%. For most proteins, there are no known molecular glues. These were used to test the ability of GlueFinder to identify possible small molecule glues in a comprehensive set of human protein dimers.

#### Small molecule screening library

All small ligands in the September 2024 Protein Data Bank (found in individual chains) were systematically extracted, and a representative library was constructed from pockets containing at least 10 amino acid residues and a minimum volume of 100 Å³. This procedure yielded 1,314 distinct ligands, which are provided in SI, LIST.ligands_library_hist.sort. The corresponding ligand–pocket PDB associations are in SI, List.templates_with_ligands_pockets_v2 which contains 94,074 structures. The distribution of the distinct ligands across pockets is highly variable, ranging from 99 instances for YCM (S- (2-amino-2-oxyethyl)-L-cysteine) to a single occurrence for ligands such as 289 (D-glycero-alpha-D-manno-heptopyranose) (63).

#### Prediction of molecular glues between E3-ligases and EGFR, HER2 and KRAS

A list of 390 human E3-ligases from NHLBI (see SI, Ligase_List.xlsx) was used to predict high confidence predictions of the complexes with EGFR, HER2 and KRAS using AF2Complex (59).

## Acknowledgments/Funding

This research was supported in part by grant GM118039 from the Division of General Medical Sciences of the National Institutes of Health. A gift from the Ovarian Cancer Institute is gratefully acknowledged. Rachel Grimley is acknowledged for her critical inputs into the manuscript. We thank Jessica Forness for polishing this manuscript and Bartosz Ilkowski for his computational support.

## Abbreviations

MGD: Molecular Glue degraders
PROTAC: proteolysis-targeting chimeras

