## Supplementary figures and images for "GlueFinder: A Data-Driven Framework for the Rational Discovery of Molecular Glues"

### Supplemental Figure S1A

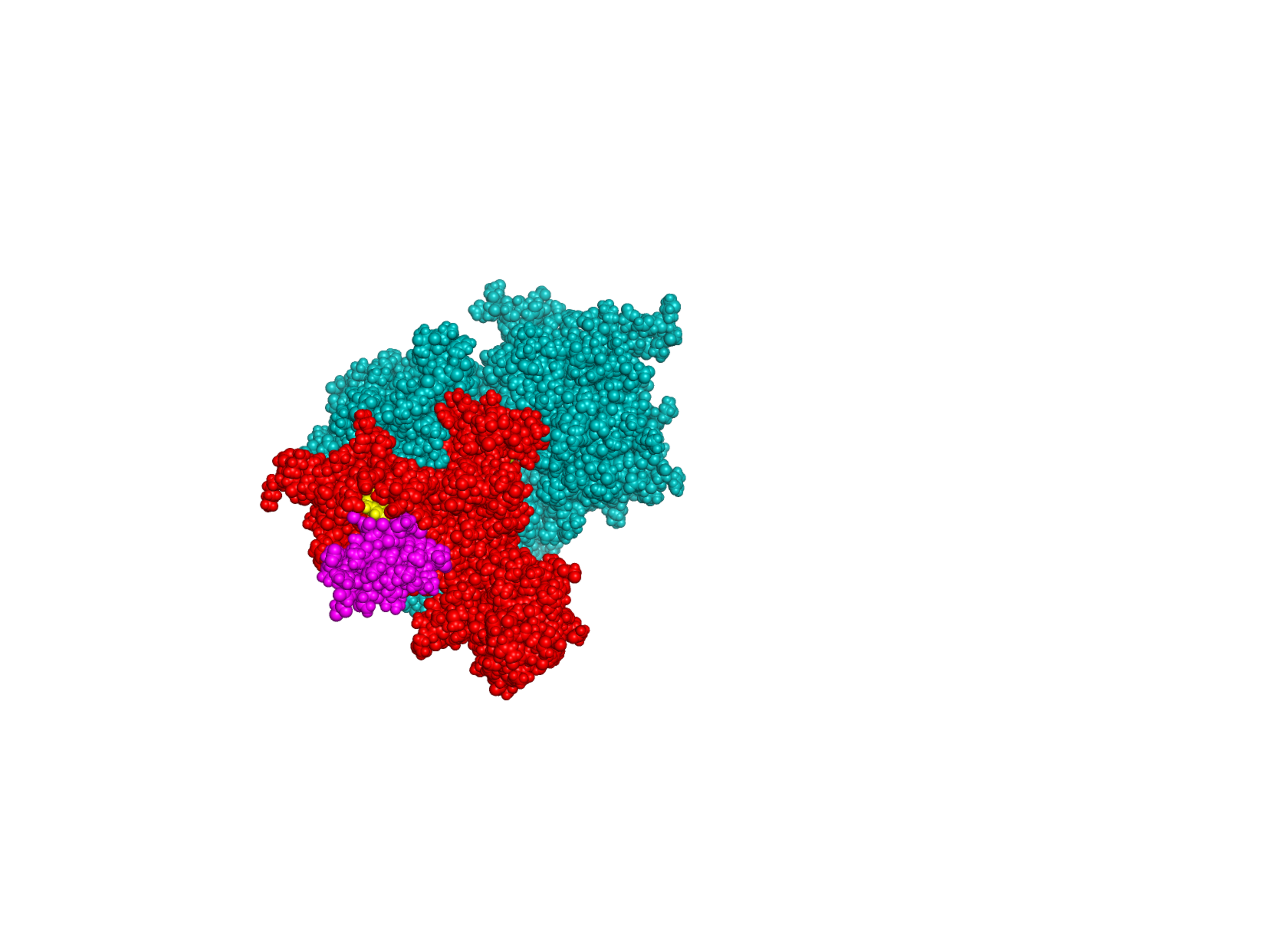

### Supplemental Figure S1B

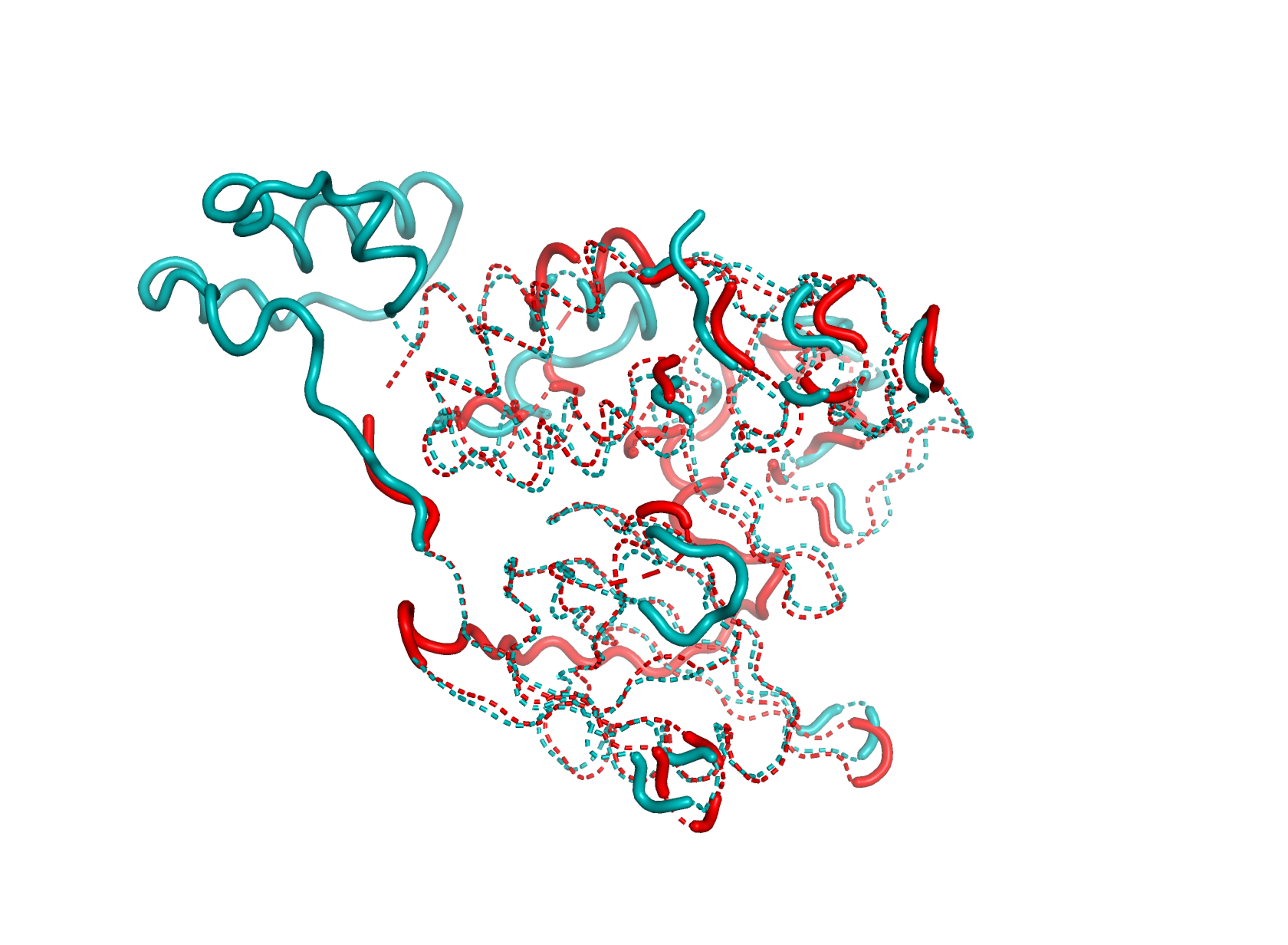

### Supplemental Figure S2

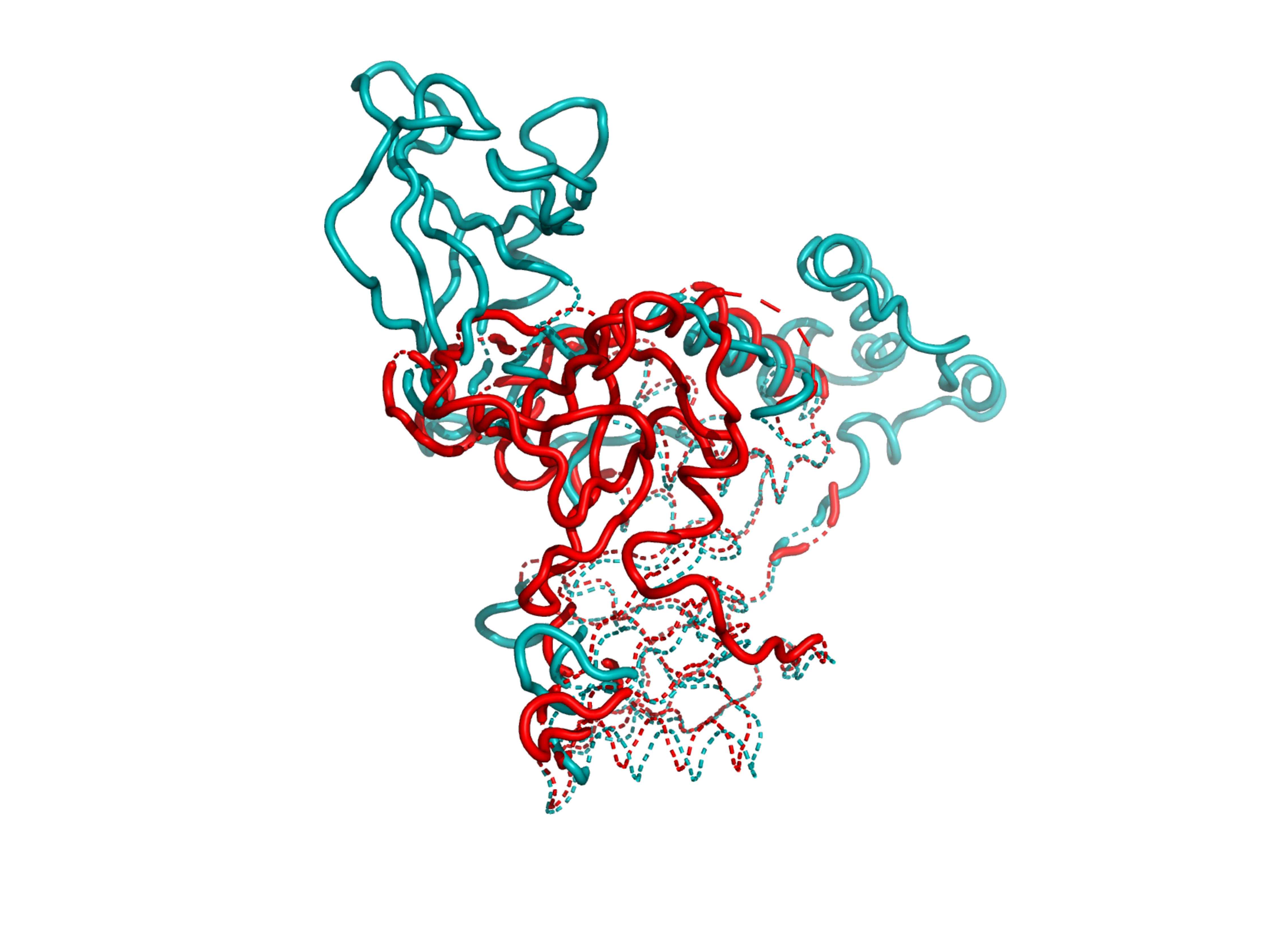

### Supplemental Figure S3

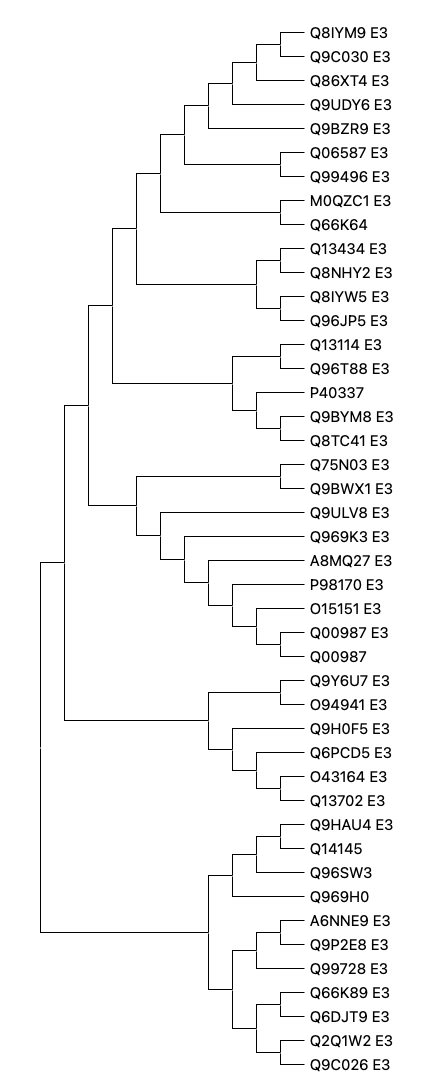
